# What is AIDS in the Amazon and the Guianas in the 90-90-90 era?

**DOI:** 10.1101/2020.03.13.990473

**Authors:** Mathieu Nacher, Antoine Adenis, Basma Guarmit, Aude Lucarelli, Denis Blanchet, Magalie Demar, Felix Djossou, Philippe Abboud, Loïc Epelboin, Pierre Couppié

## Abstract

**Introduction:** In the past decade, new diagnostic methods and strategies have appeared, HIV testing efforts and the generalization of antiretroviral therapy may have influenced the number of opportunistic diagnoses and mortality of HIV-infected patients. To test this hypothesis we compiled data on the top opportunistic infections and causes of early death in the HIV cohort of French Guiana.

**Methods:** HIV-infected persons followed in Cayenne, Kourou, and Saint Laurent du Maroni hospitals from 2010 to 2019 were studied. The annual incidence of different opportunistic infections and annual deaths were compiled. For patients with opportunistic infections we calculated the proportion of early deaths.

**Results:** At the time of analysis, among 2 459 patients, (treated and untreated) 90% had a viral load <400 copies, 91% of the patients in the cohort were on antiretroviral treatment, and 94.2% of patients on treatment for over 6 months had undetectable viral loads. Only 9% of patients had CD4 counts under 200 per mm3. Disseminated histoplasmosis clearly remained the most frequent (128 cases) opportunistic infection among HIV-infected persons followed by cerebral toxoplasmosis (63 cases) and esophageal candidiasis (41 cases). Cryptococcal meningitis was ranked 5^th^ most frequent opportunistic infection as was tuberculosis (31 cases). The trend for a sharp decline in early deaths continued (3.9% of patients).

**Conclusions:** Despite the successes of antiretrovirals, patients presenting with advanced HIV are still common and they are still at risk of dying. Improved diagnosis, and notably systematic screening with appropriate tools are still important areas of potential progress.

**Author summary:** In the past decade, new diagnostic methods and strategies have appeared, HIV testing efforts and the generalization of antiretroviral therapy may have influenced the number of opportunistic diagnoses and mortality of HIV-infected patients. To test this hypothesis we compiled data on the top opportunistic infections and causes of early death in the HIV cohort of French Guiana. HIV-infected persons followed in Cayenne, Kourou, and Saint Laurent du Maroni hospitals from 2010 to 2019 were studied. The annual number of different opportunistic infections and annual deaths were compiled. For patients with opportunistic infections we calculated the proportion of early deaths. At the time of analysis, most patients were virologically controlled and had restored immunity. However, histoplasmosis clearly remained the most frequent (128 cases) opportunistic infection among HIV-infected persons followed by cerebral toxoplasmosis (63 cases) and esophageal candidiasis (41 cases). Cryptococcal meningitis was ranked 5^th^ most frequent opportunistic infection as was tuberculosis (31 cases). The trend for a sharp decline in early deaths continued (3.9% of patients). Despite the successes of antiretroviral therapy, patients presenting with advanced HIV are still common and they are still at risk of dying. Improved diagnosis, and notably systematic screening with appropriate tools are still important areas of potential progress.

## Introduction

Nearly a decade ago we showed that the top ranking HIV-related opportunistic infectious diseases in French Guiana, a French territory in South America, were quite different from what is usually described in Europe.[1] Indeed, as immunosuppression grows, the respective frequency of AIDS-defining infections reflects the regional pathogen ecosystem. A good, knowledge of the local epidemiology is therefore crucial to guide the clinicians’ diagnostic hypotheses, their investigations and urgent therapeutic decisions. In the past decade, knowledge and awareness of regional particularities have evolved and new diagnostic methods and strategies have appeared,[2,3] HIV testing efforts may have reduced the size of the reservoir of undiagnosed HIV infections, and the generalization of antiretroviral therapy has decreased HIV incidence and improved the overall immunity of HIV cohorts.[4,5] Such evolution is likely to influence the number of diagnoses and mortality of HIV-infected patients.[6] To test this hypothesis we compiled data on the top opportunistic infections and causes of early death (1 month after an opportunistic infection) in the HIV cohort of French Guiana.

## Methods

The standards of healthcare in French Guiana are similar to those of metropolitan France.[7] All human immunodeficiency virus (HIV) patients receive free antiretroviral treatments (usually the most recent drugs) regardless of their residence status or income. Medical imagery, viral loads, CD4 counts, genotyping, antiretroviral concentration measurements, a rich diagnostic arsenal are available for routine care. The parasitology-mycology laboratory performs fungal culture. and Pasteur institute performs tuberculosis diagnoses (direct examination, PCR, and culture). The diagnosis of histoplasmosis relies on the identification of histoplasma on direct examination of samples, culture, or histopathology. Antigen testing is not available for routine care in French Guiana.

HIV-infected persons followed in Cayenne, Kourou, and Saint Laurent du Maroni hospitals between January 1, 2010 and December 31, 2019 were enrolled in the French Hospital Database for HIV (FHDH). The database includes most patients followed in French Guiana and nearly all AIDS cases. Trained technicians entered the demographic data, diagnoses, treatments, medical events, viral loads, CD4 and CD8 counts. Annual incidence of different opportunistic infections and annual deaths are compiled. For patients with opportunistic infections we calculated the proportion of early deaths (<1 month after diagnosis).

Patients included in the FHDH give informed consent for the use of their anonymized data and for the publication of anonymized results. This data collection has been approved by the Commission Nationale Informatique et Libertés (CNIL) since 1992 and the cohort has led to several international publications. The data were analyzed with R.

## Results

On December 31^st^ 2019, there were 2 459 patients in the active HIV cohort. Between 2012 and 2019 there was an average 5.5% annual increase of the cohort size. The sex ratio is 0.99 except in Western French Guiana where it was female-biased (0.62). The age group distribution was: <15 years (1.5%), 15-39 years (34%), 40-59 years (46%), and 60+ (18.5%). Overall, 33% of patients were born in Haiti (mostly in Cayenne), 18% in Suriname (Mostly in Saint Laurent du Maroni), 14% in France, 10% in Brazil, 8% in Guyana, for 12 % the information was not available, and for the remaining 5 % in a variety of countries. Among the total cohort, (treated and untreated) 90% had a viral load <400 copies, 91% of the patients in the cohort (1 749/1 921, 108 treatment naive, and 64 interruptions) were on antiretroviral treatment, and 94.2% of patients on treatment for over 6 months had undetectable viral loads (up from 78% in 2012). At the time of analysis, only 9% of patients had CD4 counts under 200 per mm3.

Figure 1 Shows the Ranking of the most frequent incident AIDS-defining infections in French Guiana between 2012 and 2019. Histoplasmosis clearly remained the most frequent opportunistic infection among HIV-infected persons followed by cerebral toxoplasmosis and esophageal candidiasis. Pneumocystosis which often ranks higher in mainland France was ranked 4^th^ and was followed by cryptococcal meningitis and tuberculosis (Fig 1).

**Fig 1.**
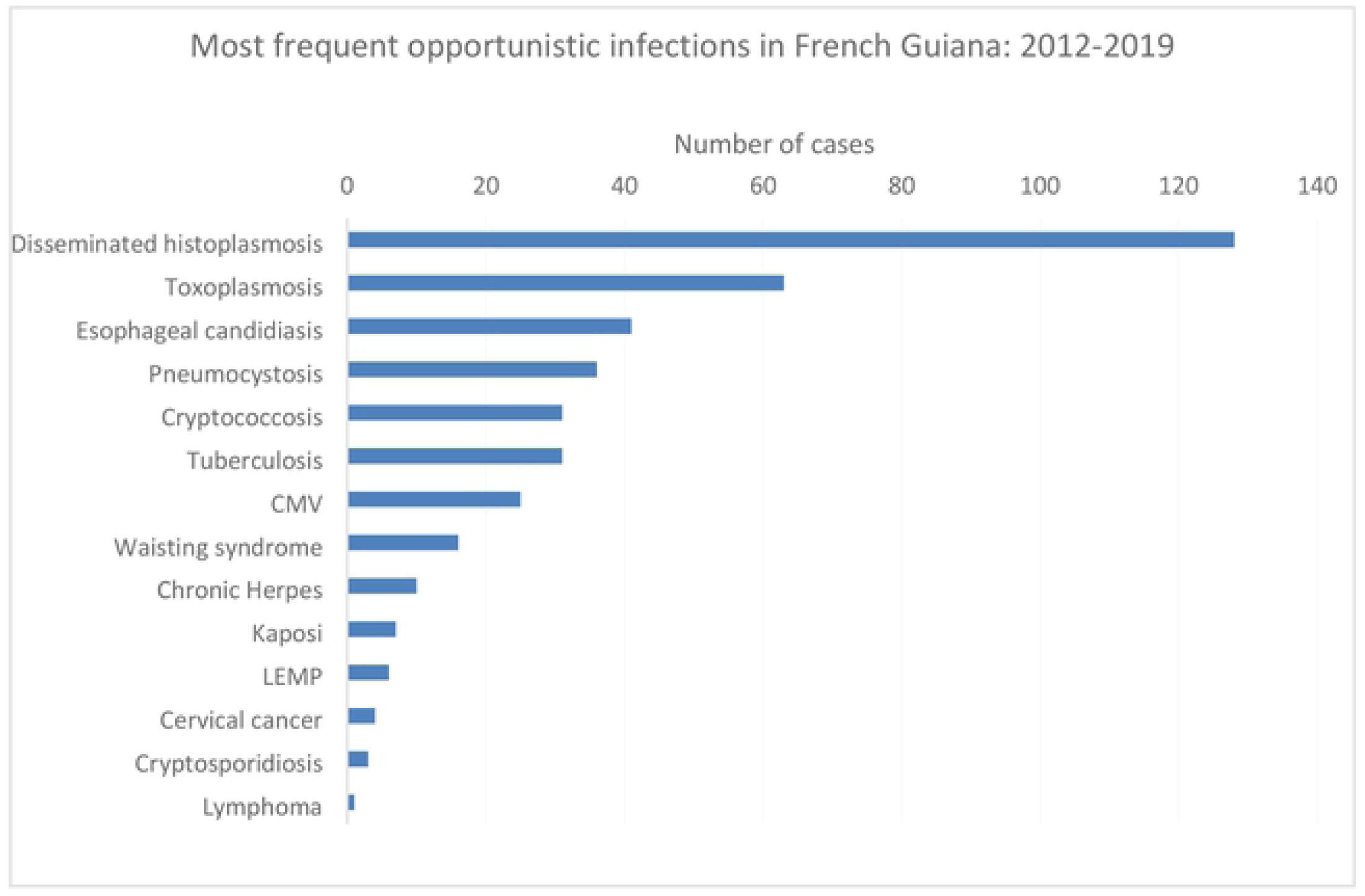
Most Frequent opportunistic infections in French Guiana 2012-2019.

Fig. 2 shows the historical evolution of disseminated histoplasmosis diagnoses and the sharp decline in early deaths (<1 month after diagnosis of histoplasmosis).

**Fig 2.**
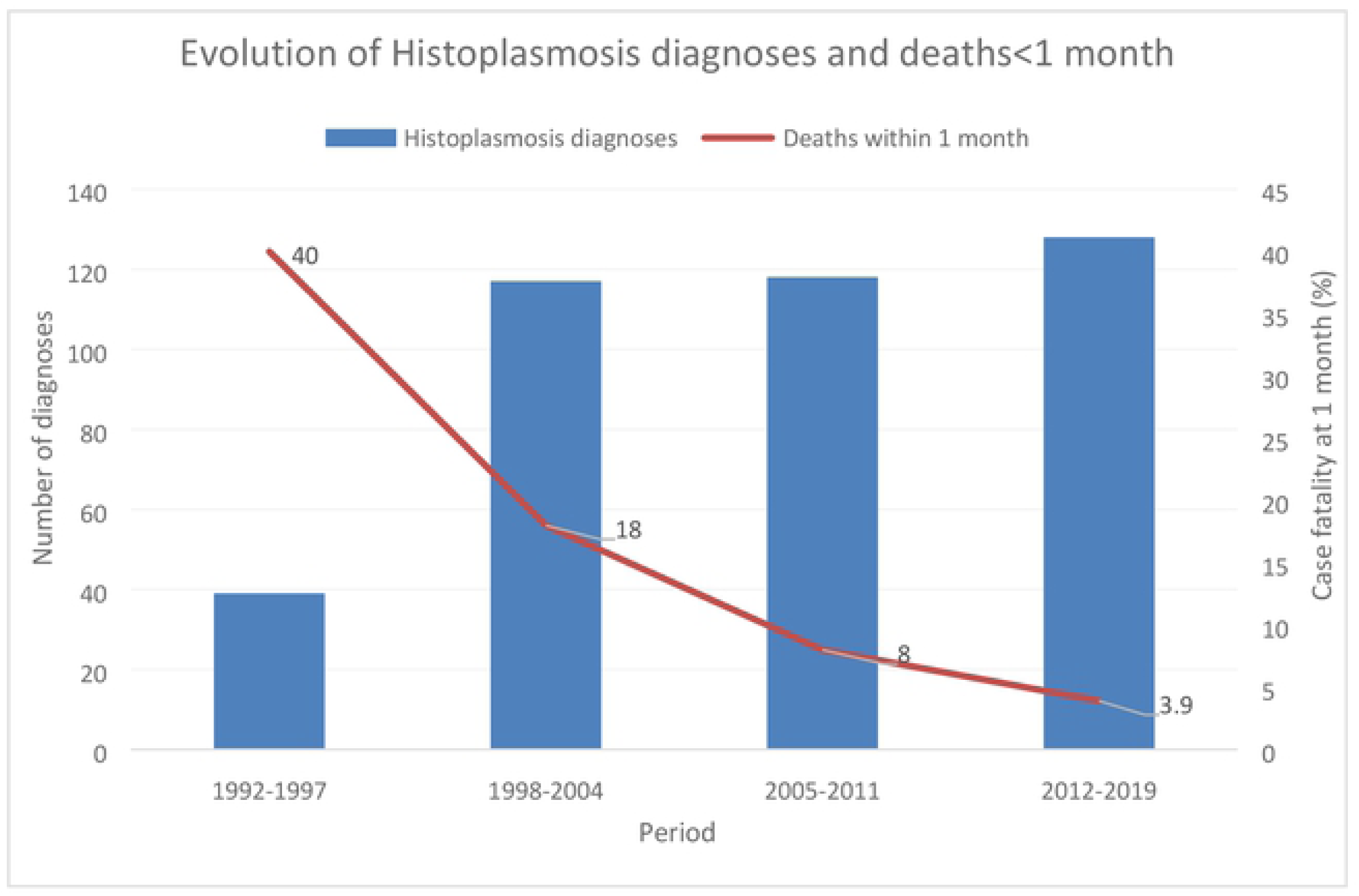
Evolution of Histoplasmosis and deaths<1 month.

## Discussion

The present results show that, despite a very high proportion of the cohort on virologically successful antiretroviral treatment, disseminated histoplasmosis remained the most frequent opportunistic infection. The case fatalities within 1-month of diagnosis, a time-frame used to reduce the likelihood of other causes of death,[6] has further decreased showing that awareness, aggressive investigations, and presumptive treatment with efficacious antifungal drugs and antiretrovirals[8] leads to good outcomes in most patients with disseminated histoplasmosis. Since our previous analysis of the main opportunistic infections[1], the implementation of systematic screening for cryptococcal antigen using the CrAg LFA may explain why cryptococcosis went from the 8^th^ most frequent to the 5^th^ most frequent opportunistic infection, a feature that surely distinguishes French Guiana from mainland France. On the contrary wasting syndromes were the 4^th^ AIDS defining event before 1996, then the 6^th^ AIDS defining event in 1996-2008 and drop to the 8^th^ position during the past decade. This suggests that improved diagnoses, and notably systematic screening with appropriate tools allowed progress in the identification of opportunistic agents, notably cryptococcus.

With the arrival of new Histoplasma antigen detection methods,[9–11] the possibility of screening for histoplasma, analogous to screening for cryptococcus[3], raises a new and important research question: should we screen all patients with less than 200 CD4 for histoplasmosis? Primary prophylaxis is in practice rarely implemented perhaps because of the fear of drug-drug interactions, antifungal resistance selection, and because physicians often expect that antiretroviral therapy will quickly restore immunity. With progress, opportunistic infections seem not so much a priority as before, and diagnosis of opportunistic infections is no longer a specific topic in French HIV expert recommendations. However, despite the successes of antiretrovirals, patients presenting with advanced infections are still not rare and even today they are still at risk of dying.[12] So for French Guiana, access to commercial histoplasma antigen detection test is important if we wish to further reduce severe forms of disseminated histoplasmosis, and detect the disease in its early stages of dissemination when its clinical presentation is not yet fully patent. For other countries, where awareness and mycological expertise is not present, where liposomal amphotericin is absent, the added value would be even greater.[13–15] This is in total alignment with the Manaus declaration in 2019 recommending access to diagnostic tests and effective treatment for all hospitals in Latin America by 2025, further emphasized by the imminent WHO/PAHO guidelines for the diagnosis and treatment of disseminated histoplasmosis in HIV patients.[16]

## References

1. Nacher M, Adenis A, Adriouch L, Dufour J, Papot E, Hanf M, et al. What is AIDS in the Amazon and the Guianas? Establishing the burden of disseminated histoplasmosis. Am J Trop Med Hyg. 2011;84: 239–40. doi:10.4269/ajtmh.2011.10-0251

2. Hillemann D, Rüsch-Gerdes S, Boehme C, Richter E. Rapid Molecular Detection of Extrapulmonary Tuberculosis by the Automated GeneXpert MTB/RIF System. J Clin Microbiol. 2011;49: 1202–1205. doi:10.1128/JCM.02268-10

3. Meya DB, Manabe YC, Castelnuovo B, Cook BA, Elbireer AM, Kambugu A, et al. Cost-effectiveness of serum cryptococcal antigen screening to prevent deaths among HIV-infected persons with a CD4+ cell count < or = 100 cells/microL who start HIV therapy in resource-limited settings. Clin Infect Dis Off Publ Infect Dis Soc Am. 2010;51: 448–455. doi:10.1086/655143

4. Nacher M. L’INFECTION VIH EN GUYANE: REVUE HISTORIQUE ET TENDANCES ACTUELLES / HIV IN FRENCH GUIANA: HISTORICAL REVIEW AND CURRENT TRENDS. 2020; 9.

5. Nacher M, Adriouch L, Huber F, Vantilcke V, Djossou F, Elenga N, et al. Modeling of the HIV epidemic and continuum of care in French Guiana. PloS One. 2018;13: e0197990. doi:10.1371/journal.pone.0197990

6. Adenis A, Nacher M, Hanf M, Vantilcke V, Boukhari R, Blachet D, et al. HIV-associated histoplasmosis early mortality and incidence trends: from neglect to priority. PLoS Negl Trop Dis. 2014;8: e3100. doi:10.1371/journal.pntd.0003100

7. Nacher M. Santé GLOBALE ET GUYANE: ÉTUDE DESCRIPTIVE ET COMPARATIVE DE QUELQUES GRANDS INDICATEURS / GLOBAL HEALTH AND FRENCH GUIANA: A DESCRIPTIVE AND COMPARATIVE STUDY OF SOME MAJOR INDICATORS. 2020; 10.

8. Nacher M, Adenis A, Blanchet D, Vantilcke V, Demar M, Basurko C, et al. Risk factors for disseminated histoplasmosis in a cohort of HIV-infected patients in French Guiana. PLoS Negl Trop Dis. 2014;8: e2638. doi:10.1371/journal.pntd.0002638

9. Nacher M, Blanchet D, Bongomin F, Chakrabarti A, Couppié P, Demar M, et al. Histoplasma capsulatum antigen detection tests as an essential diagnostic tool for patients with advanced HIV disease in low and middle income countries: A systematic review of diagnostic accuracy studies. PLoS Negl Trop Dis. 2018;12: e0006802. doi:10.1371/journal.pntd.0006802

10. Cáceres DH, Samayoa BE, Medina NG, Tobón AM, Guzmán BJ, Mercado D, et al. Multicenter Validation of Commercial Antigenuria Reagents To Diagnose Progressive Disseminated Histoplasmosis in People Living with HIV/AIDS in Two Latin American Countries. J Clin Microbiol. 2018;56. doi:10.1128/JCM.01959-17

11. Caceres DH, Scheel CM, Tobon AM, Ahlquist Cleveland A, Restrepo A, Brandt ME, et al. Validation of an enzyme-linked immunosorbent assay that detects Histoplasma capsulatum antigenuria in Colombian patients with AIDS for diagnosis and follow-up during therapy. Clin Vaccine Immunol. 2014;21: 1364–8. doi:10.1128/CVI.00101-14

12. Nacher M, Huber F, Adriouch L, Djossou F, Adenis A, Couppié P. Temporal trend of the proportion of patients presenting with advanced HIV in French Guiana: stuck on the asymptote? BMC Res Notes. 2018;11: 831. doi:10.1186/s13104-018-3944-y

13. Adenis AA, Valdes A, Cropet C, McCotter OZ, Derado G, Couppie P, et al. Burden of HIV-associated histoplasmosis compared with tuberculosis in Latin America: a modelling study. Lancet Infect Dis. 2018;18: 1150–1159. doi:10.1016/S1473-3099(18)30354-2

14. Nacher M, Adenis A, Mc Donald S, Do Socorro Mendonca Gomes M, Singh S, Lopes Lima I, et al. Disseminated histoplasmosis in HIV-infected patients in South America: a neglected killer continues on its rampage. PLoS Negl Trop Dis. 2013;7: e2319. doi:10.1371/journal.pntd.0002319

15. Caceres DH, Zuluaga A, Arango-Bustamante K, de Bedout C, Tobon AM, Restrepo A, et al. Implementation of a Training Course Increased the Diagnosis of Histoplasmosis in Colombia. Am J Trop Med Hyg. 2015;93: 662–7. doi:10.4269/ajtmh.15-0108

16. Caceres DH, Adenis A, de Souza JVB, Gomez BL, Cruz KS, Pasqualotto AC, et al. The Manaus Declaration: Current Situation of Histoplasmosis in the Americas, Report of the II Regional Meeting of the International Histoplasmosis Advocacy Group. Curr Fungal Infect Rep. 2019;13: 244–249.

